# Lagrange-NG: The next generation of Lagrange

**DOI:** 10.1101/2022.04.19.488734

**Authors:** Ben Bettisworth, Stephen A. Smith, Alexandros Stamatakis

## Abstract

Computing ancestral ranges via the Dispersion Extinction and Cladogensis (DEC) model of biogeography is characterized by an exponential number of states relative to the number of regions considered. This is because the DEC model requires computing a large matrix exponential, which typically accounts for up to 80% of overall runtime. Therefore, the kinds of biogeographical analyses that can be conducted under the DEC model are limited by the number of regions under consideration. In this work, we present a completely redesigned efficient version of the popular tool Lagrange which is up to 2.5 times faster, which we call Lagrange-NG (Next Generation). We further reduce time-to-completion by introducing a multi-grained parallelization approach, achieving a total parallel speedup of 8.5 over Lagrange on a machine with 8 cores. In order to validate the correctness of Lagrange-NG, we also introduce a novel metric on range distributions for trees in order to assess the difference between any two range inferences. Finally, Lagrange-NG exhibits substantially higher adherence to coding quality standards. It improves a respective software quality indicator as implemented in the SoftWipe tool from average (5.5; Lagrange) to high (7.8; Lagrange-NG). Lagrange-NG is freely available under GPL2.

---

The dispersal-extinction-cladogenesis (DEC) model [10] is widely used to analyze biogeographical data. However, computing likelihoods under this model is computationally challenging. This is because (i) the geographical regions are splayed into 2^*r*^ = *s* states and (ii) the computation of the respective transition matrix is in 𝒪(*s*^3^). Thus, computing a single likelihood of the DEC model requires 𝒪((2^*r*^)^3^) = 𝒪(2^3*r*^) time. In other words, the likelihood computation is exponential with respect to the number of regions under study. Therefore, the scalability of data analyses under the DEC model is limited be the number of regions, that is, only a small number of 6 to 10 regions can be analyzed in reasonable time [7].

The most expensive inference step is the computation of the transition matrix that often accounts for 80% or more of overall runtime. As in standard likelihood-based phylogenetics, the transition matrix is computed via a matrix exponential, albeit on a substantially larger matrix. Substantial research effort has been invested into finding the best way to compute the matrix exponential [8], but it still remains challenging to compute efficiently as well as accurately. Additionally, unlike in standard phylogenetics, the DEC model is non-reversible (i.e., uses a non-symmetric rate matrix), which limits the number of applicable numerical methods for computing the matrix exponential, typically to less precise ones. In the following, we present the Lagrange-NG (Lagrange-Next Generation) software, an almost complete rewrite of the popular and widely used Lagrange software by Ree et. al. [10]. As the primary challenge to computing the likelihood under the DEC model is to efficiently calculate the matrix exponential, Lagrange-NG relies on a specifically tuned version of the matrix exponential algorithm known as “scaling and squaring”[8]. Alongside the improvements to the matrix exponential, many so-called “micro-optimizations” (for example, passing function arguments by reference instead of value, using more efficient data structures to store regions, or eliminating unnecessary computation) have been implemented that further accelerate computations. We have also implemented a task-based hybrid multi-threading approach, which increases the rate of analyses by up to a factor of 8 for datasets with exceeding 200 taxa. Furthermore, we improve upon the numerical stability compared to the original software, and fix a major bug which we discovered during development. Finally, to verify that Lagrange-NG produces analogous results as the original implementation, we devised a novel method of comparing range distribution on trees, which is based on the Earth mover’s distance metric. A similar application of the Earth mover’s distance has been successfully applied to phylogenetic placement, though this method and application is distinct[3].

## 1 Software Description

Lagrange-NG constitutes an nearly complete rewrite of the original (unpublished) C++ version of Lagrange. Of the 4600 lines of code present, only 5% were present from the original code base. This redesign retains the complete functionality of Lagrange, but is computationally more efficient, and implements a parallelization of DEC calculations. Lagrange-NG implements four major improvements to Lagrange. First, it supports parallelism via a hybrid task based parallelization scheme which utilizes both coarse and fine grained parallelism. Second, it deploys more efficient numerical methods, including a custom tuned version of the matrix exponential. Third, it introduces general improvements and optimizations, that is, micro-optimizations, which individually do not notably increase efficiency, but put together yield a substantial improvement. Finally, the fourth improvement is a substantial increase in coding standards adherence and hence, software quality, as measured by the coding standards adherence evaluation tool and benchmark SoftWipe [15]. The SoftWipe score of the original Lagrange software is 5.5, while our nearly complete rewrite increases this to a score of 7.8. While the original score of 5.5 is fairly average, the new score of 7.8 places Lagrange-NG 3rd in the list of 51 scientific software tools written in C or C++ that are contained in the SoftWipe benchmark.

Importantly, during the process of improving the code quality, a potentially serious bug was discovered. In order to correct numerical instabilities, the transition matrix was normalized such that the rows summed to 1.0 after the matrix exponential computation. During normal computation, this operation will have little effect on the results. However, if the rate matrix is sufficiently ill conditioned, the computation exhibits an extreme numerical instability such that any results produced are meaningless. If the matrix is then normalized at this point, then results produced with this matrix are made to appear sensible. Therefore, any error in the computational process is hidden from the user, and the results of the computation will be perceived as plausible. Fortunately, as long as the matrix remains unnormalized, these errors are easy to detect, as several analytical conditions are no longer met (such as the rows no longer summing to 1.0). We are not aware of any approaches to recover from these errors, but at least the user is not misled into thinking that meaningless results are plausible. This normalization error is exceedingly rare, as the authors never observed it in the thousands of datasets analyzed for this paper. Despite this, the error *can* occur, and *will* by Murphy’s law. As such, we are convinced that in the event of this bug, the user should be appropriately informed. Therefore, in this case, Lagrange-NG simply fails, and alerts the user to what occurred.

Additionally, we identified and corrected a configuration error in the process of building Lagrange, where important compiler optimization options were not properly utilized. Fixing this configuration error alone increased the computational efficiency of the original Lagrange by up to 10x. While this error is easy to overlook, yet trivial to fix, we assume that many past Lagrange analyses were conducted using the unoptimized code. Nonetheless, in this work when we perform benchmarks with Lagrange, we do them with this configuration error fixed.

Lagrange-NG can be downloaded from GitHub at https://github.com/computations/lagrange-ng. To build the software, the only requirements are a C++ compiler, and CMake. Optionally, Lagrange-NG can be built with the respective system versions of the Intel Math Kernel Library (MKL) [1] and NLOpt [9, 5]. If a system version of MKL is not present, Lagrange-NG will build with OpenBLAS [2] instead.

Please see the supplementary material for a more thorough description of Lagrange-NG.

## 2 Performance

To assess the performance of Lagrange-NG relative to the original implementation, we randomly generated a large number of synthetic datasets with a varying number of regions and executed Lagrange and Lagrange-NG to record the respective runtimes. We generated 100 random datasets with either 5, 6, or 7 regions, and all had 100 taxa, to obtain a total of 300 datasets. Furthermore, we ran an additional series of parallel performance evaluations on Lagrange-NG with 8 threads assigned to coarse grained parallelization using the same datasets. The results of this performance assessment are shown in Figure 1. Additional performance tests are available in the Supplementary Material.

**Figure 1:**
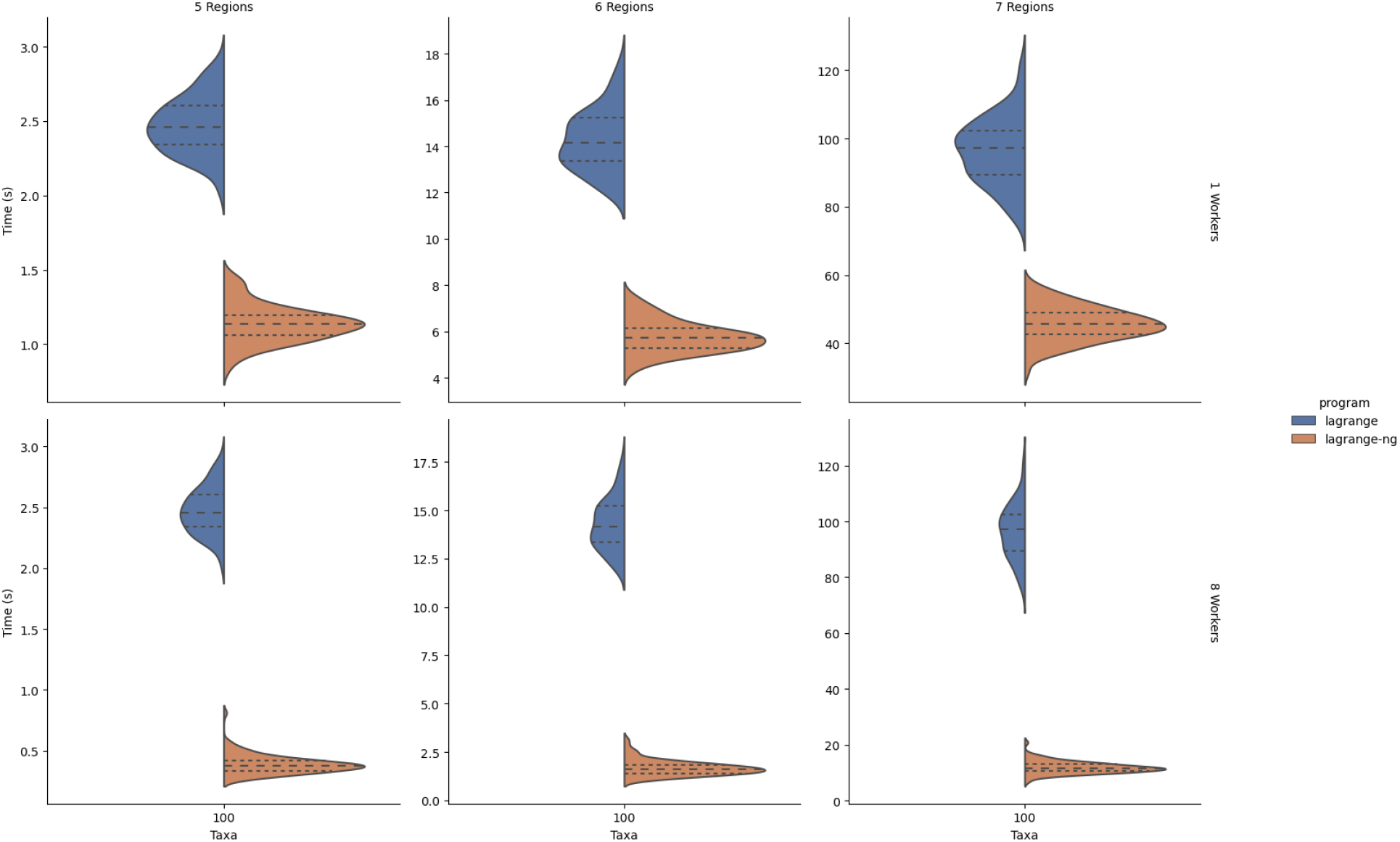
Comparison of runtimes between Lagrange (left) and Lagrange-NG (right) with sequential Lagrange-NG (top) and parallel Lagrange-NG using 8 workers (bot). Results were obtained by generating 100 random datasets. Note that the original Lagrange was not run with any multi-threading, as it does not support it. Instead, the data has been replicated for comparison’s sake. Times are in seconds. The figure was generated using seaborn [14]

## 3 Validation

Lagrange-NG re-implements core numerical routines of Lagrange. Such changes in numerical routines are often associated with difficult and subtle bugs as well as slight numerical deviations. We sought to ensure that Lagrange-NG and Lagrange produced the same results. To this end, we developed a pipeline to (i) generate random datasets, (ii) run both, Lagrange, and Lagrange-NG, and (iii) compare the results of the two programs. To compare results, we developed a measure to evaluate the distance between distributions of ancestral ranges on trees based on the Wasserstein metric [13]. We provide a summary here; further details are provided in the Supplementary material.

To compute the distance between two ancestral range distributions on a given tree, we embed the range as a vertex weight into a hypercube graph, and then use that graph to compute the Wasserstein metric between the distributions for each node. This metric is then normalized to be between 0.0 and 1.0 by dividing by the maximum total distance, and subsequently averaged over all nodes.

We ran this comparison for 100 iterations on datasets comprising 10, 50, and 100 taxa, and a number of regions between 2 and 6. This yielded 15 parameter sets, for a total of 1,500 tests.

## 4 Biological Examples

While we conducted extensive tests on simulated data, we also verify that Lagrange-NG behaves correctly on empirical datasets. To this end, we reproduced the results from a previous study on sloths from **(author?)** [12]. Additionally, we took the opportunity to reproduce the results using the tools specified in the paper as this gave us an opportunity to compare with BioGeoBEARS [7], a similar tool.

In order to reproduce results, we downloaded the supplementary data from the Dryad repository associated with the publication. To run the analyses with BioGeoBEARS and Lagrange-NG, we had to slightly modify the data. This involved correcting some taxon names so that they matched between the tree and the region data, and also removing the outgroup from the tree as there was no region data included for the outgroup. These modifications appear to be in line with what the original authors must have done, because the results from both BioGeoBEARS and Lagrange-NG match the results reported in the paper. Both BioGeoBEARS and Lagrange-NG were run with the same dataset on the same computer. Despite the fact that the original study limited the number of regions to 5, we decided to also measure Lagrange-NG’s performance with no region limit, to show that Lagrange-NG can analyse large empirical datasets without a region limit.

## 5 Results

The validation of Lagrange-NG with respect to Lagrange, was suprisingly successful, despite substantial modifications of nearly all critical code paths and numerical routines. Of the 1,500 tests, 0 produced results with differences over the tolerance of 1 × 10^−4^ when computed using our novel distance method, indicating that the results are equivalent between the two tools.

The mean sequential speedup between Lagrange-NG over Lagrange on one core for 5, 6, and 7 regions is 2.17, 2.47, and 2.11, respectively. The overall time-to-solution speedup of Lagrange-NG with 8 cores over sequential Lagrange on one core for 5, 6, and 7 regions is 6.38, 8.5, and 8.11, respectively.

In addition, Lagarange-NG is substantially faster than BioGeoBEARS. On the empirical dataset, Lagrange-NG computed the result in about .5 minutes using 8 cores, while BioGeoBEARS required about 14 minutes to analyze the data using 80 cores. BioGeoBEARS and Lagrange-NG inferred different optima for model parameters, with BioGeoBEARS achieving a slightly better log-likelihood score (−216.127 vs -224.396). It is unclear if these likelihoods are directly comparable. Nonetheless, this does not affect the respective qualitative results as BioGeoBEARS and Lagrange-NG agree on the most likely distribution for every node.

The analysis of this dataset with no region limit using Lagrange-NG produced similar results to the analysis with the 5 region limit, albeit with a better likelihood (−217.023). The time for this analysis was also .5 minutes using 8 cores.

## 6 Discussion

We have shown that computation of likelihood-based biogeographical models can be greatly accelerated without sacrificing result quality. An 8 fold increase in speed over the original implementation, and a 28 fold increase in speed over BioGeoBEARS represents a step forward, especially when taking the time complexity of the matrix exponential into account. Additionally, we retain this speed even on datasets with a large number of regions and no region limit, enabling for more fine grained as well as exploratory analyses of biogeographical data.

Readers might wonder why the execution time for the analysis of the empirical dataset with a maximum number of regions is the same as the analysis without the maximum number of regions. Ostensibly, a smaller number of regions should lead to a faster execution, but the equal runtime between the two analysis seems to counter this. However, there is a core function in the likelihood computation that depends on the specific states. When there is a region limit placed on the analysis, the fine-grained execution flow of this function becomes unpredictable for the CPU, which leads to missed branch predictions (branch prediction tries to predict the outcome of conditional statements in the code to accelerate computations) by the hardware. In turn this induces a slower likelihood calculation. Therefore, the speed gain from the region limit seems to be, in this case, exactly matched by the speed loss from incorrect branch predicitons. Lagrange-NG is optimized for the no region limit case, and as such implements trade-offs which favor this case. Thus, analyses with a region limit should be avoided if the only reason for them is performance.

However, the parallel efficiency of 0.5 is rather sub-optimal. Yet, as detailed in the Supplementary Material, the parallel efficiency increases with increasing taxon and region numbers (strong scaling). This means that, as datasets become more taxon rich or as the size of the transition matrix grows, and run-times increase, Lagrange-NG will utilize its parallel computational resources more efficiently.

Future work on Lagrange-NG includes extending the range of models that can be computed by the DIVA/DIVALIKE and BAYAREA family of models [6, 11]. Additionally, the current models can be further optimized in three areas, although we expect unspectacular performance improvements.

We produced a version of Lagrange-NG that utilized GPU acceleration for the matrix computations. Unfortunately, this method failed to produce acceptable speedups even for large datasets (10-11 regions). This is in line with the performance results of previous attempts to accelerate likelihood computations for phlyogenetic tree inference on GPUs [4]. The fundamental difficulty is that the tree-based nature of the computation that induces a decreasing degree of parallelism as we approach the root, leaves many computational units starved for work, as is the case with the existing CPU-based course-grained parallization. It is possible that further development would produce better results, but we believe that by the time that datasets become large enough to observe large speedups, the analysis will simply be infeasible due to the exponential nature of the problem.

For remaining improvements to the computational efficiency of Lagrange-NG, the first is to further refine the matrix exponential routine. While we settled on a particular implementation with good performance, there are other, more exotic methods such as Krylov-type methods which entirely avoid computating *e*^*A*^ and might yield better performance. Finally, one could further refine the work allocation for the coarse grained parallelization approach. The current method of assigning tasks is straight-forward, and can be improved upon by becoming aware of which nodes are “most blocking” of other tasks. It might be possible to devise an algorithm that can minimize the “task starved” period of computation, either as a clever assignment method, or as a so-called “work stealing” scheme.

## 7 Availability

The software, tools, and data used for this paper are available online at https://github.com/computations/lagrange-ng.

## 8 Supplementary Material

Additional material describing the operation of Lagrange-NG, as well as additional scaling experiments can be found in the Supplemental Material.

