## Supplemental Material for "Lagrange-NG: The next generation of Lagrange"

### LAGRANGE-NG: THE NEXT GENERATION OF THE DEC MODEL

A PREPRINT

**Ben Bettisworth**
Computational Molecular Evolution
Heidelberg Institute for Theoretical Studies
Heidelberg, Germany


**Stephen A. Smith**
Ecology and Evolutionary Biology
University of Michigan
Ann Arbor, Michigan


**Alexandros Stamatakis**
Computational Molecular Evolution
Heidelberg Institute for Theoretical Studies
Heidelberg, Germany


April 19, 2022

#### 1 Background

Lagrange-NG implements the DEC (Dispersion Extinction and Cladogensis) model of geographic range evolution [8]. A geographic range, in this context, describes the broadly defined distribution of the habitat of a particular species. The evolution of this range is assumed to follow the phylogeny, or the biological evolution of a species or clade. The DEC model takes, as a minimum, a phylogenetic tree, and a set of regions. The phylogenetic tree is assumed to be the true phylogeny of the included species, and the regions are the generalized areas of potential habitation for the species in question. The DEC model constructs a list of states based on the valid set of regions that a particular species could inhabit. With these components, the DEC model constructs a transition matrix between states using two parameters, an extinction parameter and a dispersion parameter. Using this transition matrix, a likelihood of the model parameters can be computed and used to optimize the model parameters. Once the optimal model parameters have been found, the most likely ancestral ranges can be found by computing the model "backwards".

Computation of the likelihood of a particular set of parameters under the DEC model proceeds in a fashion similar to the standard Felsenstein pruning algorithm [4]. In this algorithm, the computation starts from the tips and moves towards the root, storing intermediate results in buffers called conditional likelihood vectors. Please see [13] for a more detailed explanation, including a detailed discussion about the savings involved with such a scheme. What is relevant for this discussion is that the Felsenstein pruning algorithm avoids excess computation by noticing that, at certain points in the computation of a likelihood on a tree, the only relevant quantity is the likelihood *conditioned on the current state*.

#### 2 Methods and Algorithms

Lagrange-NG utilizes a task based parallelization scheme in which each node of the tree is assigned as a task. In order to compute the results for a generic node of the tree results for its two children must first have been computed. This involves computing

1. The right and left instantaneous rate matrices: $Q_r$, $Q_l$,
2. The right and left transition probability matrices: $P_r = e^{Q_r t_r}$ and $P_l = e^{Q_l t_l}$ respectively,
3. The result of the Markov process along the left and right branches: $w_r = P_r v_r$ and $w_l = P_l v_l$ respectively,
4. The weighted combination of $w_r$ and $w_l$, $v_t$.

Together, these operations make up a single task for a worker. However, as it can be seen above, the task can be further subdivided into smaller parts, which we will call operations. For the purposes of this paper, we will label each of the operations as

1. Make Rate Matrix Operation,
2. Expm Operation,
3. Dispersion Operation and,
4. Split Operation.

In order to more easily support parallel computation on other platforms, such as GPUs, we have separated the operations from the memory buffers which are required to store the intermediate results needed for likelihood computation. For example, in Figure 2 we show a generic node and its associated operations. For each operation, there are a set of indices which indicate the location of the assigned memory buffer. Additionally, they store the last execution clock point, details and purpose of which are discussed later.

Operations are typically fast enough that they cannot be separated into parallel tasks. However, if the model parameters and branch lengths are appropriate, the results of these operations can be shared between two tasks. For example, consider the tree topology in Figure 1. Here, two of the branches have the same branch length, 1.0. If they also share a rate matrix (which they almost always will), then the result of the two matrix exponential operations will be identical. Therefore, the likelihood computation on this tree can be accelerated by computing $e^{Qt}$ only once, and saving the result. An operation can be shared when the model parameters (the extinction and dispersion rates) and the branch lengths are the same between two branches.

In order to avoid such redundant computations, Lagrange-NG can share operations between tasks. However, when computing with multiple threads, this introduces the possibility of conducting computations with inconsistent values from dependant operations when performing successive evaluations of the likelihood with model parameters that are altered by optimization routines.

To avoid race conditions, I.E. cases when one thread is reading data that is not ready, we use a clock based method to enforce a partial ordering on the computation of operations. Readers familiar with vector clocks will recognize this as a vector clock with the number of distributed elements equal to one. After the evaluation of each operation, a "time" is recorded in the evaluated operation, and the clock incremented. This time is not a true time, but instead a virtual time that is incremented every time an operation is completed. To determine if an operation can be carried out, a thread only needs to check if the clocks of its immediate dependant operations show a larger value. If this is the case, then the dependant operations have already been evaluated, and the operation can be performed on consistent input data.

By carefully dividing the tasks into operations, merging identical operations between tasks, and enforcing a partial order on the computation of operations, we can implement an effective task-based parallelization scheme. However, this scheme does have an unavoidable bottleneck. Since the dependencies for the operations is based on the topology of the tree, and the tree has fewer branches near the root than near the tips, the threads will necessarily be work starved near the end of a round of computation. Therefore, our parallel efficiency is limited, and in fact dependant on, the topology of the tree.

##### 2.1 Coarse and Fine Grained Parallelization

Most linear algebra libraries offer some sort of BLAS level parallelization, which is what Lagrange-NG uses for its "fine-grained" parallelism. However, in early testing we found that the parallel efficiency of this mode of parallelism was quite poor for our purposes, often making the run slower for a moderate number of threads and a small number of regions (for example, 6 threads and 5 regions). So, we elected to a implement a coarse-grained parallelization method, which parallelized over the available operations. To distinguish these two modes of parallelism, we named the coarse-grained threads "workers" and the fine-grained parallelism "threads per workers".

#### 3 Comparing Distributions on Trees

When comparing the results of Lagrange-NG and Lagrange, it is difficult to assess if the numerical results are "close enough" to be the same. By way of example, scientific software will typically, instead of asking if two floating point numbers are exactly identical to one another, ask if they are closer than some small number called $\epsilon$. This is due to the limitations in the IEEE 754 floating point number format, which is how real numbers are encoded in nearly all modern computers. A more complete explanation of this phenomenon is outside the scope of this paper, and is extensively

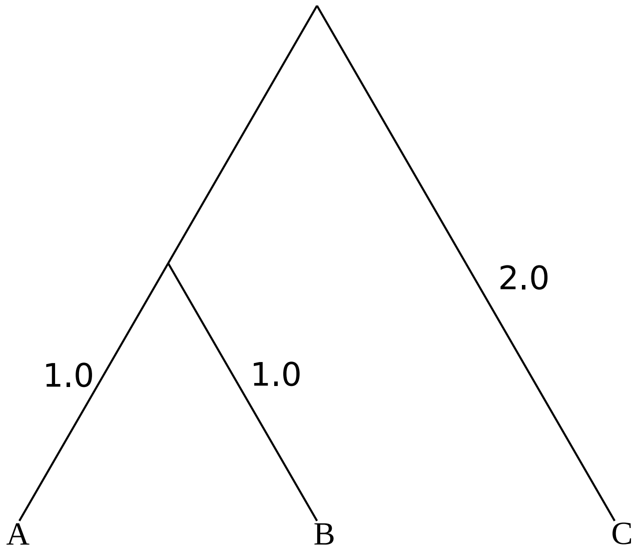


Figure 1: A simple ultrametric tree.

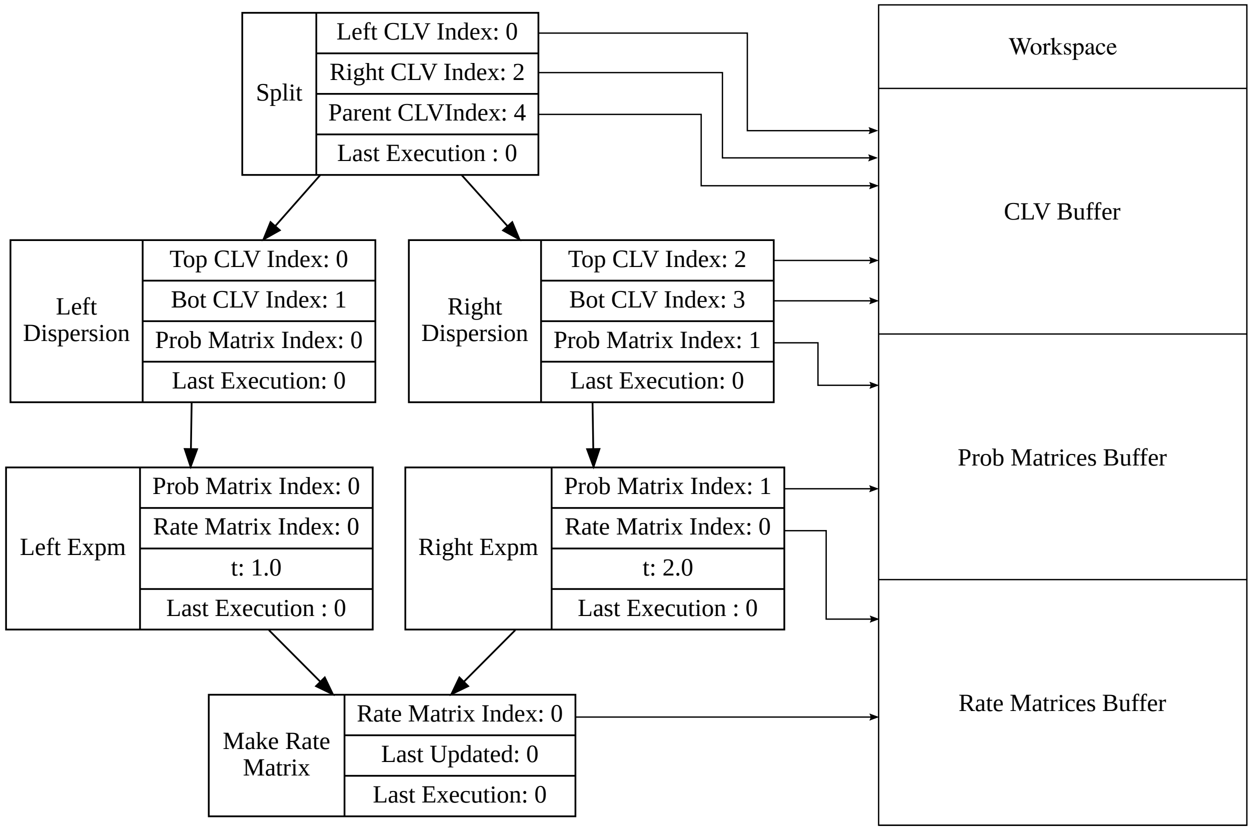


Figure 2: An example set of operations for an generic unspecified node.

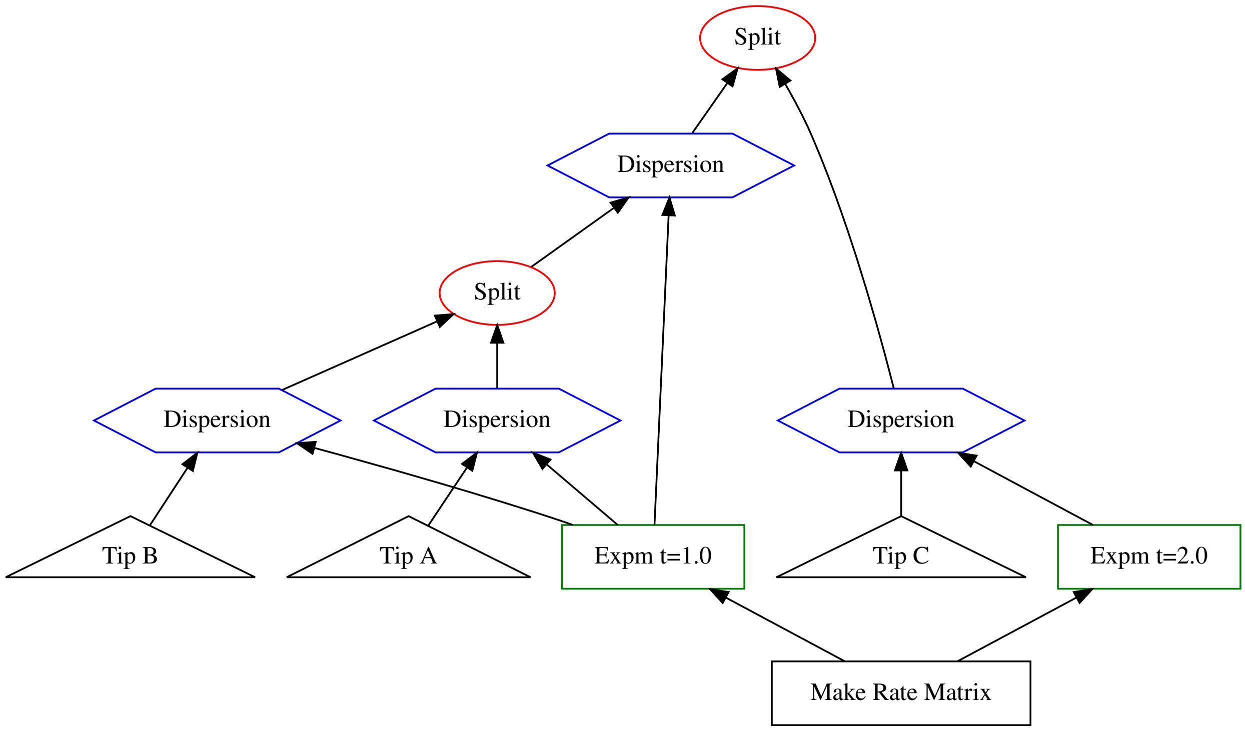


Figure 3: Tree in Figure 1 decomposed into the operations used to compute the likelihood.

discussed in any textbook on numerical computing, but for our purposes it is sufficient to say that differences in the order of associative mathematical operations will generally yield different results.

Lagrange-NG does not escape this limitation of the IEEE 754 standard. So, in order to appropriately compare results, we must take this into account. One can examine, by eye, each distribution individually and compare the distributions by hand, but this is time consuming, subjective, and error prone. Therefore, it is desirable to construct a metric between the node base probability distributions on trees in order to automatically compare the results.

Consider an example, where we have two sets of ancestral range distributions computed by differing methods using the same tree. Let us call those distributions $d_1$ and $d_2$. The distribution for node $n$ is then represented by the notation $d_i(n)$. Since the tree topology is identical for the two distributions, we can match the node level distributions in a one-to-one mapping between the two sets of distributions. We will index the individual elements of the distribution either by the list of regions names or the binary notation for regions. For instance, if we have a distribution over regions $A, B$, and $C$, then the entry for the distribution $AB$ for node $n$ would be indexed as $d_1(n, AB)$. Equivalently, we can use a binary notation to write $d_1(n, 110)$. In this case $A$ stands for the most significant bit, $B$ the second most significant bit, and $C$ the least significant bit.

In this example, the first approach would be to treat $d_1(n)$ and $d_2(n)$ as vectors, and simply compute the cosine distance between distributions. This will indeed produce a metric, but it has some undesirable properties. For example, suppose we have a distribution over 5 regions: $A, B, C, D$, and $E$ where $d_1(n, AB) = 1.0$. If we use the cosine distance to compute the distance between $d_1(n)$ and $d_2(n)$ where $d_2(n, ABC) = 1.0$, then the resulting distance will be 1.0, as the vectors are orthogonal. However, this is the same distance as if we had $d_2(n, CDE) = 1.0$ instead which is also orthogonal to $d_1(n)$. But, the prediction $AB$ is much closer to $ABC$ than $CDE$. In the first case, the predicted ranges differ by only *one* region, whereas in the second case, the predicted ranges differ by *every* region.

Since the cosine distance does not account for the available transitions between states in the DEC model, we should pick a distance that is aware of these transitions. To accomplish this, we first embed the two distributions into a hypercube graph. A hypercube is graph with $2^n$ nodes and each node is connected to $n$ other nodes (see Figure 4 for an example). Importantly, the edges of a hypercube graph correspond to the valid transitions between states in the DEC model[1].

[1]Some readers might have noticed in Figure 4 that the edges don't distinguish a direction of the edges, which means that transitions *out* of the extinct state are valid. While conceptually this is a problem, for the purposes of computing a distance, it will

Once the distributions are embedded in a hypercube graph, we can compute the distance between the distributions as the "amount of effort required to turn one distribution into the other". This is the Wasserstein metric, also known as the Earth mover's distance, and is what we base our distance off of. Suppose we have the distributions $D1$ and $D2$ from Figure 5. In order to transform $D1$ into $D2$, we need to find a way to move $0.25$ "earth" from node `10` to node `01`. Two possible example transformations can be seen in Figure 6. While the distance computation in Figure 6 is straightforward, in general, finding the minimum transformation distance requires the use of an optimization routine.

To find the minimum distance, we formulate the problem as a linear programming problem. Specifically, we solve

$$\begin{aligned} &\min_x \textstyle\sum_i x_i \\ &\text{such that } Ax = b \\ &x_i \geq 0 \end{aligned} \tag{1}$$

Where $A$ is the constraint matrix induced by the hypercube, and $b = d_1(n) - d_2(n)$, that is, the difference between the two distributions. If we have a distribution with $s$ states then we can produce $A$ by creating $s - 1$ rows and $s(2^s - 1)$ columns. The rows represent the nodes of the graph, and the columns represent the edges of the graph, split in two for each direction of flow. The entries of $A$ are defined as:

$$A(n, e) = \begin{cases} 1 & \text{If } e \text{ points to } n \\ -1 & \text{If } e \text{ points away from } n \\ 0 & \text{Otherwise.} \end{cases} \tag{2}$$

Please note that there are only $n - 1$ rows. This is because the final row can be expressed as a linear combination of the previous rows, and will therefore induce no additional constraints to the problem. Additionally, we choose to suppress edges leading away from the extinct state, to be consistent with the model. By suppressing these edges, we remove $s$ columns from the matrix as well.

As an example, the distance between the distributions in Figure 5 can be computed with the matrix

$$A = \begin{pmatrix} 1 & 0 & 1 & 0 & 0 & 0 \\ -1 & -1 & 0 & 0 & 1 & 0 \\ 0 & 0 & -1 & -1 & 0 & 1 \end{pmatrix}$$

and the vector

$$b = \begin{pmatrix} 0 \\ 0.25 \\ 0.25 \end{pmatrix}.$$

To actually compute the solution, we can turn to any of a number of linear programming solvers. Finally, in order to normalize the distance, we divide the result by the maximum possible path length, which is simply the number of *regions*.

#### 4 Selecting the Best Linear Algebra Library

In the course of the development of Lagrange-NG, we tested and benchmarked several different linear algebra libraries. The tested libraries are: OpenBLAS[3], Blaze[7], Eigen[5], and MKL[2]. Of these, the best performing libraries are MKL, followed by OpenBLAS. However, these two libraries are within 10% performance of each other for our purposes.

In addition to testing various linear algebra libraries, we also tested several methods of computing the matrix exponential. For all libraries, we tested the scaling and squaring method that eventually became the final method. For Blaze and Eigen, an eigenvalue based method was also tested. We implemented these two version in Blaze and Eigen only because of their rich support for linear algebra algorithms, beyond what is available in BLAS and LAPACK.

Of the methods tested, scaling and squaring based implementations performed the best for both speed and stability. The eigenvector based methods regularly had stability problems, and therefore were ineligible for final implementation.

Additionally, two GPU based versions of Lagrange-NG were tested, one using cuBLAS[1], and the other using MAGMA[9]. Both implementations used the scaling and squaring method that we had determined to be the best in CPU based code. However, these version performed poorly when compared to the CPU versions, typically being at least 2x slower for the data sizes we were testing with.

not affect the results, as taking the path through the extinction state is equivalent to taking any other path of equal distance. Due to the nature of the hypercube, this second path must exist.

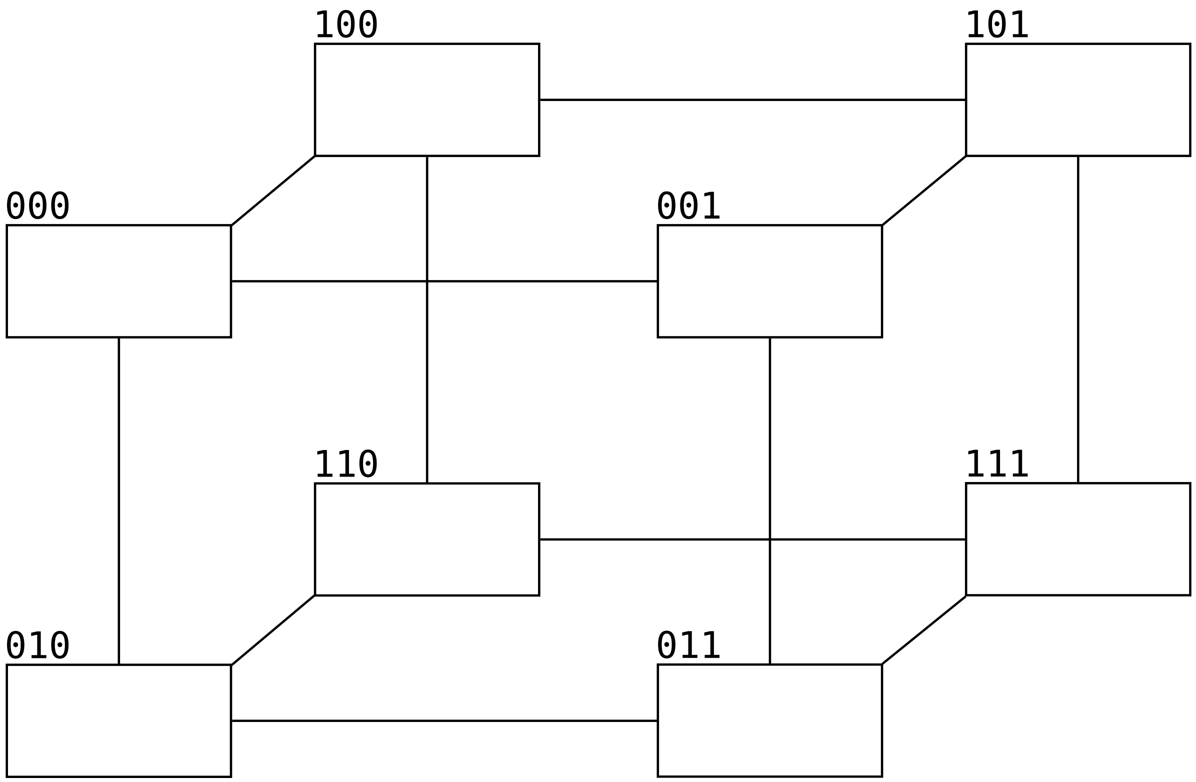


Figure 4: An example of a 3-dimensional hypercube with associated region names in binary notation.

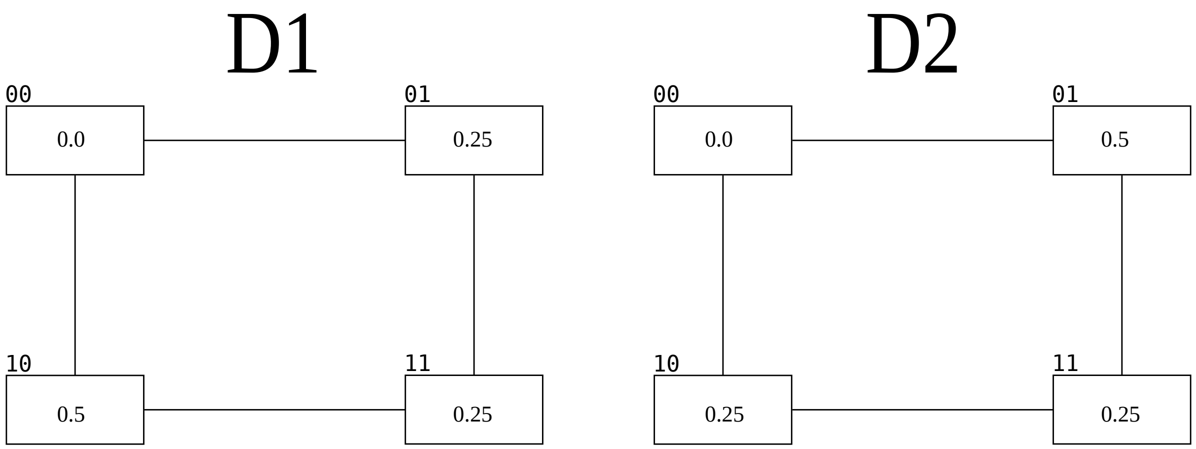


Figure 5: Two example distributions, displayed as 2d-hypercubes (squares).

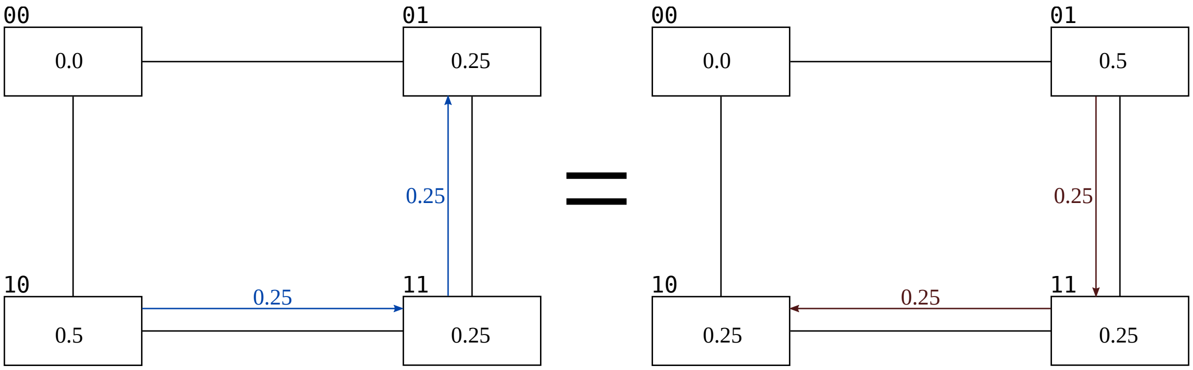


Figure 6: Distributions from Figure 5 with the Wasserstein metric applied. On the left, we choose to move 0.25 units of "mass" from node `10` to node `01`. This requires 2 transitions, shown in blue, for a total distance of 0.5. On the right, we choose to transfer mass from node `01` to node `10`, which yields the same result.

### 5 Experiments

Tests and tooling for experiments in this paper are written in Python 3 [10], with additional support from Numpy [6], SciPy [11], and seaborn [12].

#### 5.1 Investigating the Optimal Threading Configuration

Because Lagrange-NG implements both fine- and coarse-grained parallelization, we need to investigate the optimal threading configuration, that is, the number of fine-grained threads per task to be used, for a given dataset. To this end, we generated datasets with 50 or 100 taxa and 5, 6, 7, or 8 regions. This yielded a total of 8 distinct parameter sets. As the fine grained parallization scheme is nested inside of coarse grained threads, total number of threads used by Lagrange-NG is the product of fine grained threads and coarse grained threads. So, in addition to these 8 parameter sets, we also generated the 6 threading configurations with a total number of 32 threads each, as 32 can be factored with: 1 and 32; 2 and 16; 4 and 8. Since order matters, the configuration of 2 workers and 16 threads per worker is different than the configuration of 16 workers and 2 threads per worker. For each of these parameter set and threading configurations, we ran 100 trials and recorded the times. The results from these runs can be seen in Figure 7.

#### 5.2 Determining the parallel efficiency of Lagrange-NG

Given the results from the previous experiment to determine the optimal threading configuration, we choose to determine the parallel efficiency of Lagrange-NG using only workers. This is to say, we only increased the number of threads allocated to the coarse grained tasks. To this end we tested Lagrange-NG with 1, 4, 8, 16, and 32 threads by generating 100 datasets for each threading configuration. We did this with datasets with 6 regions and 100 and 500 taxa. We computed the mean of the execution times for the runs with a single thread, and used this value to compute the respective speedups.

### 6 Discussion

Regarding the optimal threading configuration, Figure 7 shows that allocating all the available cores to coarse-grained threads is normally optimal. Occasionally, allocating 2 fine threads per worker is slightly faster. This is slightly surprising, and might indicate that the linear algebra library used has potential issues with lock contention. If this is the case, then changing the library might improve results. However, there are still sequential parts of the likelihood computation which do not benefit from the fine grained parallelization, so this will have improvement a limit induced by Amdahl's law.

The threading efficiency of Lagrange-NG's coarse grained parallelization ranges between 0.74 and 0.14. We feel that this is expected, as the method of parallelization is based on the tree topology. Children nodes must be evaluated before parent nodes can be evaluated, which leads to a dependencies which prevent perfect parallel efficiency from being achieved. This means that, for every likelihood evaluation, there is a phase in the computation where there is less work available than threads. In this case the excess threads idle. However, we expect that as the number of taxa grows, the

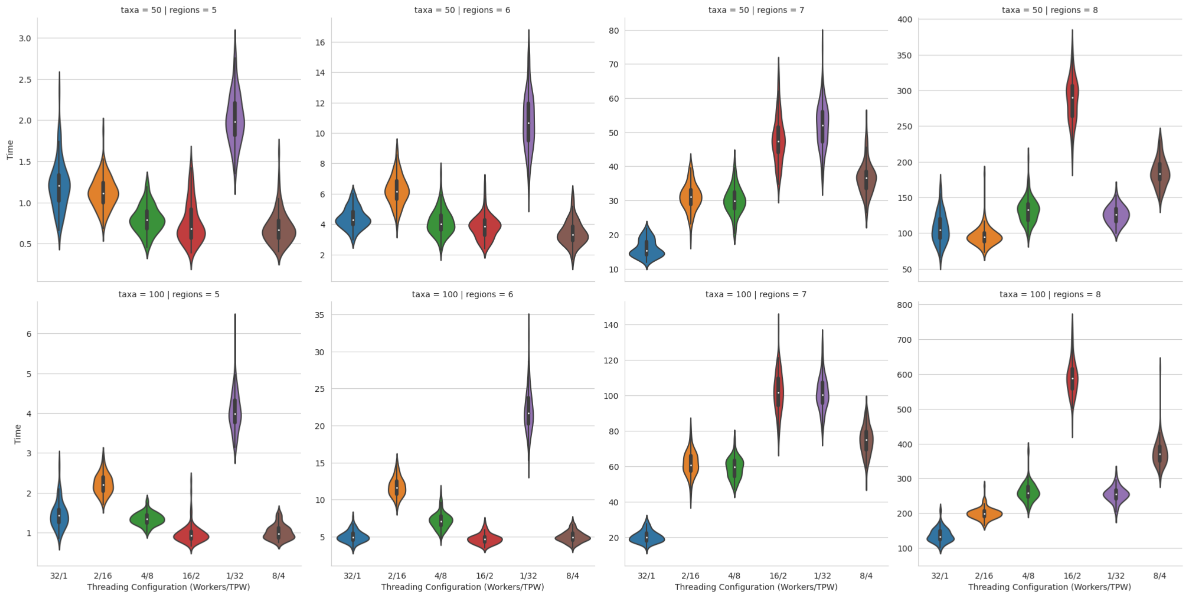


Figure 7: Plot of the threading configurations on various dataset sizes. 100 datasets were generated for each Taxa, Region, and Threading Configuration. Each dataset was generated randomly, similar to how datasets are constructed in the rest of this work.

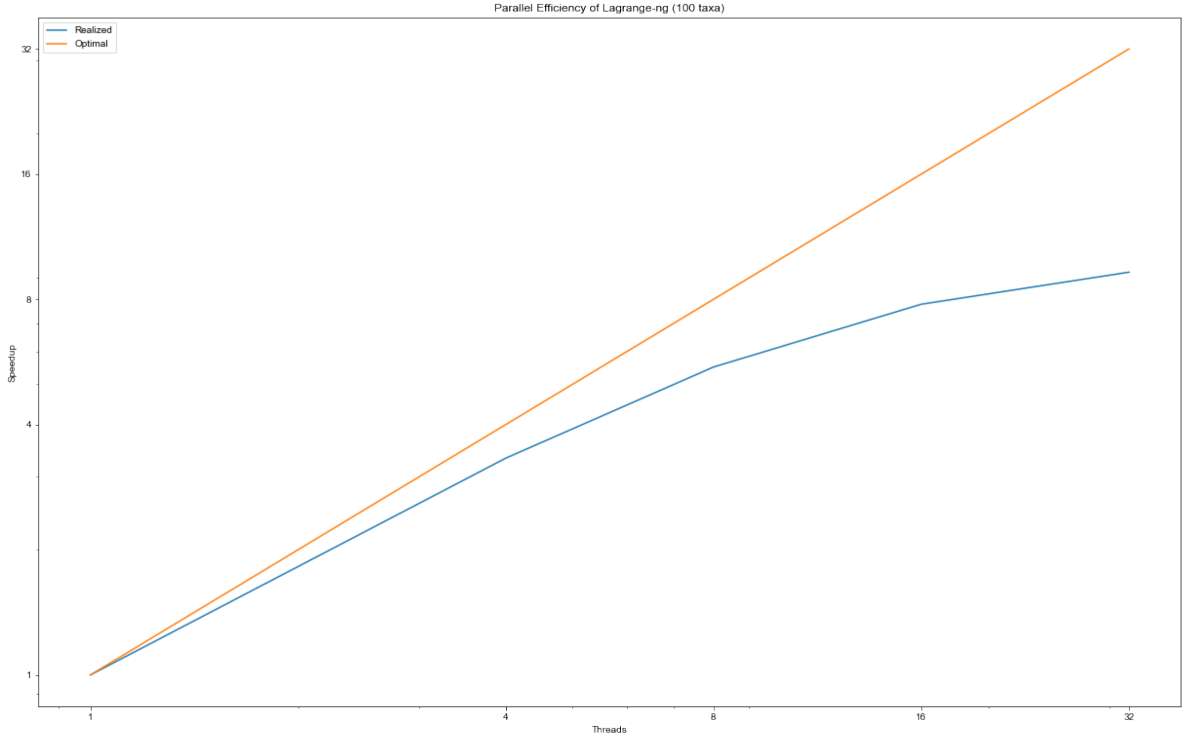


Figure 8: Parallel efficiency plot for a datasets with 100 taxa and 6 regions. Please notice the log-log scaling. The actual values plotted are 3.0, 4.1, 4.7, 4.8 for 4, 8, 16, 32 threads, respectively. The ratio of the realized speedup to the optimal speedup is 0.742873, 0.513696, 0.295814, 0.149044 for 4, 8, 16 and 32 threads respectively.

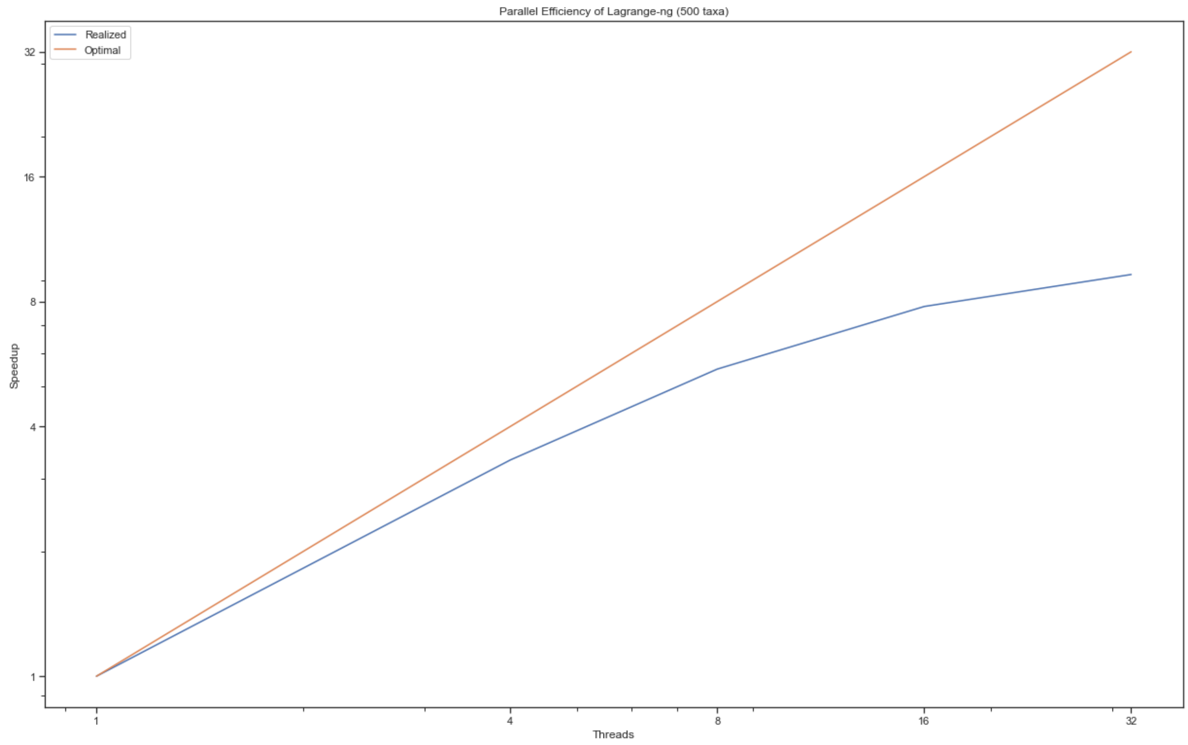


Figure 9: Parallel efficiency plot for a datasets with 500 taxa and 6 regions. Please notice the log-log scaling. The actual values plotted are 3.3, 5.5, 7., 9.3 for 4, 8, 16, and 32 threads, respectively. The ratio of the realized speedup to the optimal speedup is 0.83, 0.69, 0.49, 0.29 for 4, 8, 16, and 32 threads, respectively.

efficiency of this method should increase, as the proportion of time spent in this "work starved" phase near the root of the tree is lower. Results from the threading configuration experiments suggest that efficiency can be improved further on some datasets by allocating 2 threads per worker.

#### References


[1] cuBLAS. Archive Location: CUDA API References.

[2] Developer Reference for Intel® oneAPI Math Kernel Library - C.

[3] OpenBLAS : An optimized BLAS library.

[9] TOMOV, S., DONGARRA, J., VOLKOV, V., AND DEMMEL, J. Magma library version 0.1, 2009.

[10] VAN ROSSUM, G., AND DRAKE, F. L. *Python 3 Reference Manual*. CreateSpace, Scotts Valley, CA, 2009.

[11] VIRTANEN, P., GOMMERS, R., OLIPHANT, T. E., HABERLAND, M., REDDY, T., COURNAPEAU, D., BUROVSKI, E., PETERSON, P., WECKESSER, W., BRIGHT, J., VAN DER WALT, S. J., BRETT, M., WILSON,

J., MILLMAN, K. J., MAYOROV, N., NELSON, A. R. J., JONES, E., KERN, R., LARSON, E., CAREY, C. J., POLAT, İ., FENG, Y., MOORE, E. W., VANDERPLAS, J., LAXALDE, D., PERKTOLD, J., CIMRMAN, R., HENRIKSEN, I., QUINTERO, E. A., HARRIS, C. R., ARCHIBALD, A. M., RIBEIRO, A. H., PEDREGOSA, F., VAN MULBREGT, P., AND SCIPY 1.0 CONTRIBUTORS. SciPy 1.0: Fundamental Algorithms for Scientific Computing in Python. *Nature Methods 17* (2020), 261–272.

[12] WASKOM, M. L. seaborn: statistical data visualization. *Journal of Open Source Software 6*, 60 (2021), 3021.

[13] YANG, Z., ET AL. *Computational molecular evolution*, vol. 284. Oxford University Press Oxford, 2006.
